# Daily activity rhythms, sleep, and pregnancy are fundamentally related in the Pacific beetle mimic cockroach, *Diploptera punctata*

**DOI:** 10.1101/2025.02.27.640076

**Authors:** Ronja Frigard, Oluwaseun M. Ajayi, Gabrielle LeFevre, Lilian C. Ezemuoka, Sinead English, Joshua B. Benoit

## Abstract

Sleep and pregnancy are contentious bedfellows; sleep disorders and disturbances are associated with adverse pregnancy outcomes, although much is still unknown about this relationship. Sleep and pregnancy have been studied in many models, but most focus heavily on mammals. However, pregnancy is ubiquitous across the animal kingdom – a hallmark of convergent evolution; similarly sleep is a shared feature across diverse species. Here, we present an ideal model in which to study the dynamics between sleep and pregnancy in invertebrates. The Pacific beetle mimic cockroach, *Diploptera punctata*, is a viviparous cockroach species that uses milk proteins to nourish its young with a broodsac over a three month pregnancy. However, little is known about the relationship between this unique reproductive biology and daily rhythms of activity and sleep. We established that *D. punctata* displayed a peak in activity shortly following sunset, with males significantly more active than females. When scavenging behavior was examined, males and non-pregnant females emerged more often and traveled further from a shelter compared to pregnant females, suggesting reduced risk-taking behavior in late pregnancy. Chronic disturbance of sleep during pregnancy negatively impacted embryo development by increasing gestational duration and decreasing the transcription of milk proteins. These findings indicate that sleep is key to embryo development and that pregnancy has a significant impact on the daily rhythms of activity in *Diploptera punctata*. More broadly, we present a tractable invertebrate model for understanding the relationship between sleep and pregnancy, which will aid in the exploration of the poorly understood interface between these two ubiquitous and highly conserved traits.

## Introduction

Sleep is a ubiquitous adaptation in animals, from single celled flagellates to mammals. The benefits of sleep include neural regeneration and pruning important to memory formation and maintenance, increased immune strength, decreased exposure to predators, and conservation of energy (Keene and Duboue, 2018). Animals that experience sleep deprivation show serious defects and with eventual death (Everson et al., 1989), demonstrating that sleep is key to biological functions (Brinkman et al., 2024). Understanding sleep is critical to understanding many key features of life: it is pervasive, with diverse and critical functions, and is highly variable across and within species, and even among life stages. Interestingly, there are strong interactions between sleep and another common evolutionary feature; live birth.

While it is believed that sleep evolved divergently, with a common ancient ancestor as the source of the behavior (Anafi et al., 2019), live birth is one of the most common convergent evolutionary strategies for animals, with viviparity documented in 21 of 34 phyla including squamates, mammals, fish, and invertebrates (Ostrovsky et al. 2016). Insects are a particular hotbed for the evolution of viviparity, with over 65 instances of independent evolution of live birth documented in flies alone, and countless others amongst other insect orders (Kalinka, 2015; Meier et al., 1999). In humans (Sharma et al., 2016) and other mammals (Pires et al., 2021), pregnancy is associated with an increased incidence of sleep disorders and disruption, and these sleep disorders are associated with adverse pregnancy outcomes (Palagini et al., 2014). These interactions are heavily studied in humans; however, many assume that the systems in play are too complex to study in invertebrates. In fact, although animals without a central nervous system tend to have sleep that is controlled metabolically rather than neuronally (Anafi et al., 2019), sleep can still be characterized in lower orders. Sleep and its functions in insects have been investigated and can be quantified in a variety of ways, such as posturally, arousal threshold, and periods of immobility (Ajayi et al., 2023a; Helfrich-Förster, 2018). To our knowledge, no study has investigated the direct relationship between sleep and pregnancy in invertebrates, which leaves open the questions: do pregnant individuals follow similar sleep and activity patterns to their non-pregnant counterparts, and how important is sleep for successful pregnancy?

Here, we address these questions in an insect model of viviparity, the Pacific beetle mimic cockroach, *Diploptera punctata*, a member of the family *Blaberidae*. Cockroaches in this family, which also includes *Blaptica dubia* and *Gromphadorhina portentosa,* exhibit a range of reproductive strategies from oviparous to ovoviviparous and viviparous (Rugg and Rose, 1984). Matrotrophic viviparity, direct continual nourishment of the embryo within the reproductive tract, is famously characteristic of placental mammals but is also found in some reptiles and fish as well as tsetse flies, and *Diploptera punctata* (Benoit et al., 2015; Kalinka, 2015; Marchal et al., 2013; Roberts et al., 2016). In this way, *Diploptera punctata* are unique among cockroaches. Their eggs lack a yolk, and young are nourished inside the brood sac by milk proteins until birth (Marchal et al., 2013; Williford et al., 2004). As in mammals, this reproductive mechanism trades a greater number of young for increased survival of fewer, more developed progeny, with juvenile *Diploptera* developing within the protected brood sac at nutritional expense to the mother (Greven et al., 2014).

Genomic analyses of *Diploptera* indicate that the evolutionary transition from oviparity to viviparity requires changes in the mother to allow for gestating embryos, including physical structures to house the embryos (Fouks et al., 2023). Similar genomic changes are observed in convergent instances of live birth among many insects (Fouks et al., 2023). These adaptations include circulatory and urogenital remodeling, likely to promote gas and nutrient exchange with the brood sac, and cuticle restructuring to allow for growth (Benoit et al., 2015; Fouks et al., 2023; Marchal et al., 2013). Along with these physical changes, genomic changes in immune and endocrine systems likely support the growing embryos (Fouks et al., 2023).

One of the largest and most obvious shifts must be the supply of nutrients to the developing embryos by the mother. In *Diploptera punctata*, mothers nourish their offspring with glandular secretions of a crystalline milk-like protein into the brood pouch (Banerjee et al., 2016; Williford et al., 2004). The process of making milk is energetically expensive for viviparous insects (Attardo et al., 2012; Benoit et al., 2015, 2014; Jennings et al., 2020). While other viviparous arthropods (e.g. *Glossina*) feed on blood to support juvenile production, which is high in protein and lipids, *Diploptera* feed on far less nutrient-dense sources, such as plant proteins and other general organic materials. With this assumed increased nutritional requirement, foraging behavior must increase or maternal metabolism must slow during embryo development. Sleep deprivation is known to increase metabolic efficiency in *D. punctata* (Stephenson et al., 2007), but previous studies used only male cockroaches, and thus the effect of sleep deprivation on pregnancy is unknown. In fact, *Diploptera* have been shown to eat more regularly during pregnancy than their non-pregnant counterparts, though a significant difference in the amount of food consumed was not shown (Greven et al., 2014), implying that there is another solution to meet the embryos’ nutritional requirements. One possible answer may lie in changes to behavior and activity levels.

Many cockroach species, such as *Periplanera americana* (Lipton and Sutherland, 1970), and *Blatta orientalis* (Gunn, 1940), are nocturnal and vespertine, with a peak in activity during the first half of the night. However in *P. americana*, adult females showed activity rhythms related to ootheca deposition rather than light cycle. Like most members of *Blaberidae*, *Diploptera* are known to be nocturnal, showing the highest activity levels during scotophase (dark phase), and low activity levels during photophase (light phase) (Stephenson et al., 2007).

However, deeper study of daily rhythms, particularly in females of the species, have been neglected to this point. The unique biological aspects of pregnancy in *D. punctata* suggest that there are likely changes in sleep and daily rhythms as pregnancy progresses.

In this study, behavioral and rhythmic changes related to pregnancy are investigated in *D. punctata*, and the effects of chronic sleep deprivation on pregnant females are demonstrated. Our results reveal that there are links between daily rhythm changes and pregnancy stage, indicating decreased boldness for exploration in pregnant females during key scavenging periods. Furthermore, chronic sleep deprivation shows detrimental effects to vital pregnancy functions, such as decreases in milk protein transcription levels and gestational period elongation. This study highlights that sleep and activity patterns are shifted during pregnancy in this cockroach and that sleep periods are critical to allow for optimal milk production and minimizing the duration of pregnancy cycles.

## Materials and Methods

### Diploptera punctata maintenance

*Diploptera punctata* colonies were kept at 25°C under a constant 12 h:12 h light:dark (L/D) cycle at 70-80% relative humidity. Clear bins with air holes (Pioneer Plastics) were provided with several paper towel roll tubes for shelter, and the diet consisted of fish food (Tetramin) and dog food (Old Roy). Each colony contained about 300-400 individuals (mixed sexes and stages), and three colonies were incubated to provide the necessary sample sizes.

Sex and pregnancy status of individuals were determined based on standards established in Jennings et al. (2019). Three groups were differentiated; male, non-pregnant female, and pregnant female. Pregnancy status was based on visible pregnancy, and as ‘non-pregnant’ females had been housed with males, this group likely contained both non-pregnant females and those early in pregnancy, likely from 0-2 weeks. Likewise, ‘pregnant’ females selected for trials were in late stages of pregnancy, likely 7-10 weeks after eggs began to develop in the brood sac. This selection was accomplished based on pregnancy characteristics defined as an extended abdomen beyond the length of the wing tips and visible white banding on the abdomen. Any females who gave birth during the activity tracking trials were removed from the study as this causes tracking of both the mothers and 1st instar progeny.

### Assessment of basic parameters of diurnal activity : Single monitor trials

Basic activity and sleep levels were assessed using the Locomotion Activity Monitor (LAM) (LAM25-Trikinetics), which we have used to monitor changes in other insect systems (Ajayi et al., 2024; Bailey et al., 2024). The LAM uses infrared beams to monitor activity by logging each beam break (9 beams). Individuals were placed into *Drosophila* tubes containing standard fruit fly media (Benoit et al., 2020; Polak et al., 2023) and provided with a paste of fish food (Tetramin) and a wet sponge at the other end of the tube. Tubes were placed into the monitor (beam in the center of the tube) and sex/pregnancy status groups were alternated to ensure no placement bias (Fig. 1a). A dimming system (Lutron) was set, beginning at 17:00 and dimming by ten percent every six minutes until full darkness at 18:00. The inverse, beginning at 05:00 and brightening by the same gradient until 06:00 am, was set to create a full day-night cycle with sunrise and sunset periods. The same gradient was instituted in the colony incubators at this time to allow acclimation for several days prior to the beginning of the trial. Activity was monitored over a 7 day period, and the resulting data was analyzed using Rethomics (Geissmann et al., 2019).

**Fig. 1.**
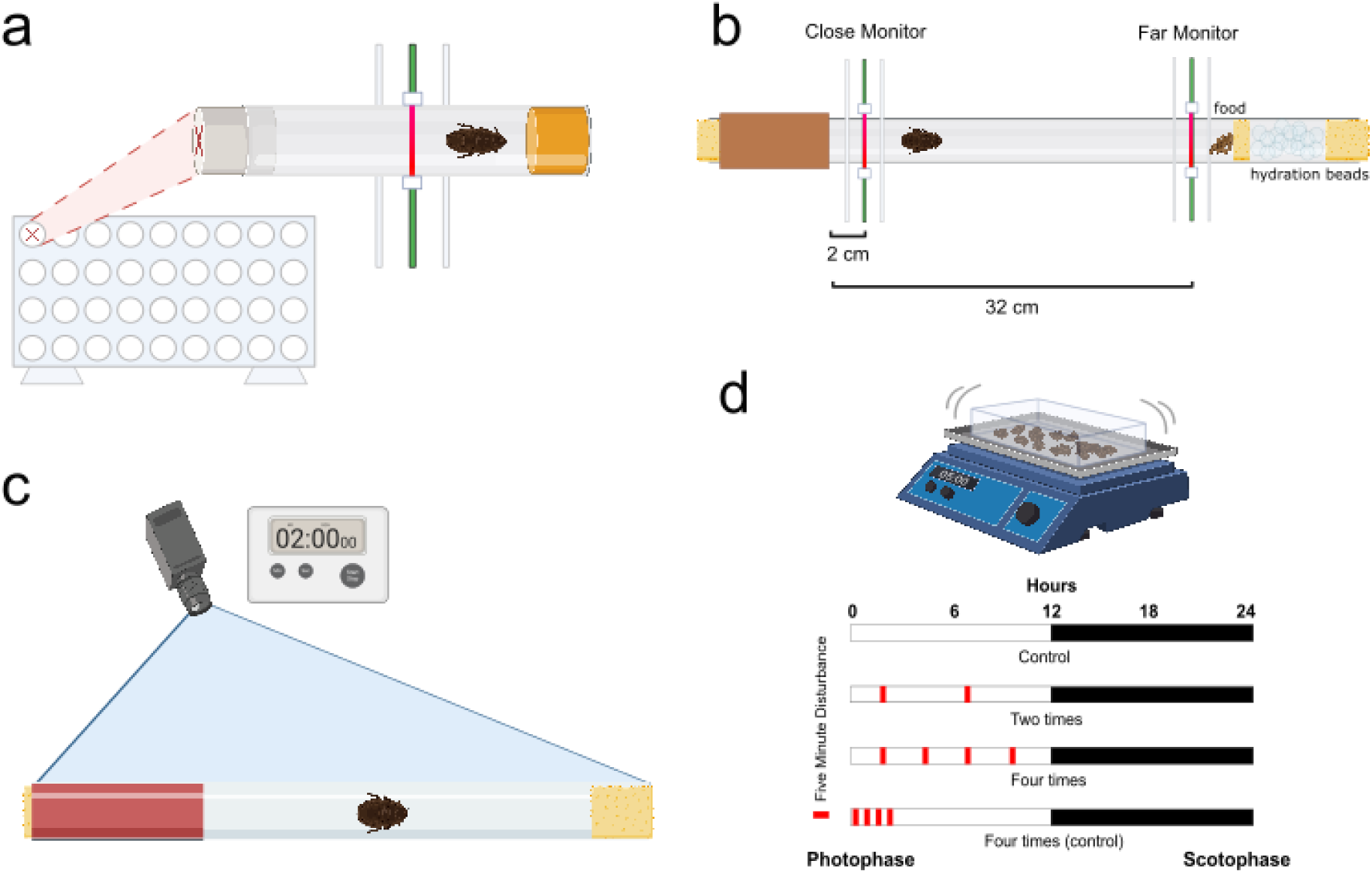
Diagram of Assay Setups. **a. Diagram of setup for Single Monitor Trials.** The Locomotor Activity Monitor (LAM25 Trikinetics) is shown from the front, with a callout showing the setup of individual *Drosophila* tubes from the side. A single monitor holds 30 tubes with cockroaches, each associated with multiple infrared beam detectors. Activity is logged when the beam is broken by movement of the *Diploptera.* **b. Diagram of setup for Dual Monitor Trials.** These trials had a similar setup as the Single Monitor Trials, but included a harborage and access to food and hydration to encourage exploration. They also utilized two LAMs instead of one, monitoring 32 46 cm tubes at distances 2cm and 32 cm from the mouth of the harborage (left). **c. Diagram of setup for Video Trials.** A similar tube as in the Dual Monitor Trials was used, but no food or hydration beads were used, allowing for a longer arena. In addition, the cardboard harborage from the Dual Monitor Trials is replaced with a red film harborage to allow investigators to view cockroaches while inside the harborage. **d. Diagram of setup for Chronic Disturbance Trials.** A shaker was used to disturb colonies for 5 minutes at a time. This was done either twice (5 and 10 hours into the photophase period), four times (2.5, 5, 7.5 and 10 hours into the photophase), or condensed into the first 2 hours of the photophase period. Created in BioRender. Frigard, R. (2024) BioRender.com/x73l781, BioRender.com/o34j999, BioRender.com/j49x931.

Across almost all species, sleep is typically characterised by decreased response time, arousal, and postural changes (Keene and Duboue, 2018). In cockroaches specifically, posture, lack of movement, and waking states can be identified as associated with cockroach sleep-liek stats (Helfrich-Förster, 2018). Based on these trends, sleep in *Diploptera punctata* has been determined at the threshold of 30 minutes with no movement (Ajayi et al., 2023b, 2022; Helfrich-Förster, 2018). Therefore, a decrease in activity can be correlated to an increase in sleep, yet there are differences, and data can be categorised into sleep and waking/activity minutes. Periods (6 hours) at the beginning and end of the 7-day observation window were trimmed to remove any influence by investigator movement and allow for an acclimation period. Dead individuals or other disturbed trials (power outage, fire alarm, etc.) were removed from subsequent analyses, resulting in an *n* of 80 individuals per group. The 24 hour data frame was split into two phases (06:00-17:59 = Day, 18:00-05:59 = Night) based on the lighting schedule.

### Basic assessment of cockroach emergence from harborages: dual monitor trials

Based on differences in activity and sleep patterns among sex/pregnancy status groups, a dual activity tracking system was used to more accurately assess willingness of cockroaches to emerge from the harborage and to determine how many of those trips resulted in exploration of the entire tube. Dual-tube LAM25 assays were conducted using the groups above, but were modified to assess cockroach emergence and movement from a harborage. A clear 46 cm tube of ⅞ inch in diameter was plugged on both ends with damp sponges and 70% RH Humidity Beads (CheapHumidors) to ensure hydration. A harborage was created on one end of the tube using paper towel rolls fastened outside of the tube, and on the other end, fish food (Tetramin Goldfish flakes) slurry was placed to encourage scavenging. One ‘close’ monitor was placed at 2 cm from the mouth of the harborage, and another ‘far’ monitor was placed at 2 cm before the food at the terminal end of the tube, leaving an empty 30 cm length of tube between the two monitors (Fig. 1b). Activity was thus monitored both at the ‘close’ emergence site and at the ‘far’ scavenging site. The experiment used a 12:12 L:D cycle, with dimming and brightening occurring on a timer to simulate natural sunrise and sunset periods. Sunrise began at 06:00 and brightened by 5 percent every three minutes until full light at 06:30, sunset began at 18:00 and dimmed by 5 percent every three minutes until total darkness at 18:30 pm. Activity levels were assessed using methods similar to those based on a single monitoring system, and the day was split into several phases (05:31-08:00: sunrise, 08:01-17:30: day, 17:31-20:00: sunset, 20:01-05:30: night) based on the lighting schedule for comparison of activity in relation to sex and pregnancy status. For males *n* = 28, for non-pregnant females *n* = 33, and for pregnant females *n* = 31. This discrepancy is a result of any individuals which died or gave birth during the trial being removed from the dataset.

### Detailed analysis of harborage use and emergence at sunset: video tracking

Video tracking was conducted focusing on a two-hour period encompassing sunset with the same 46 cm tubes as above. For this trial duration, hydration and nutritional sources were not provided other than a single moist sponge placed at each end of the tube. The harborage for these trials consisted of a half cylinder of cardboard to provide a ‘floor’ with the tube covered in a transparent red film to provide dark conditions. A camera (GoPro, model SP1M1) was mounted approximately two feet above the tubes, with full visibility, and a dimming system controlled a full spectrum LED light (Fig. 1c) The light dimmed as above, beginning at 18:00 and dimming by 5 percent every three minutes until it reached 0%. A red bulb was also used, allowing investigators to view the movement of the cockroaches via video in relative darkness. Individuals were placed into the tubes several hours prior to the beginning of recording to acclimate. Recordings were taken for 2 hours: 45 minutes of full light, 30 minutes of dimming, and 45 minutes of full darkness. Using the DLTDV8 system (Hedrick, 2008), a deep learning program was trained to analyze the video footage and track specimen movement. Training videos were from both light and dark periods to ensure accuracy. Points generated were analyzed and separated by hand into individual emergences before averages of depth, number, and duration were evaluated. For Number, *n* = 22 males, 21 non-pregnant females, and 19 pregnant females. Difference in group size is due to the removal of certain individuals due to unusable videos (truncated, disturbed, poor video quality, etc). Duration of a single emergence was defined as the time from when a cockroach fully left the harborage until the time when the individual returned. Depth refers to the distance traveled from the mouth of the harborage towards the far end of the tube. For Duration, *n* = 100 males, 29 non-pregnant females, and 20 pregnant females were used. For Depth, *n* = 100 for males, 29 non-pregnant females, and 20 pregnant females. Differences in n for Duration and Depth groups is due to different total numbers of emergences per group.

### Pregnancy length and progeny number following sleep deprivation: chronic disturbance trials

In order to establish the specific gestational length, juvenile cockroaches were reared individually to ensure virgin females and allowed to mate by giving multiple males access to nearly emerged females. Females can then be observed over a several month period to record pregnancy length and outcome. Following successful mating, pregnant individuals were kept in separate bins (11.6 x 10.2 x 3.8 cm, Pioneer plastics) with standard rearing conditions and a small harborage as described above. Disturbances occurred either not at all (control), twice (5 and 10 hours into the photophase period), four times (2.5, 5, 7.5 and 10 hours into the photophase), or condensed into the first 2 hours of the photophase period to allow the cockroaches a 10 hour period undisturbed in the photophase (secondary control). The secondary control group was included to ensure that effects on pregnancy were due to sleep disturbance as opposed to mechanical damage or other side effects of disturbance. ‘Disturbance’ was achieved by holding colonies on a shaking plate for five minutes.

Pregnancy duration was defined as the time from mating until birth. Individuals that did not give birth after 110 days were removed from the analyses as abortion of the first cohort of eggs likely occurred. This cutoff was based on previous data which defines the maximum pregnancy duration at 100 days (Engelmann, 1959). We also measured the total number of progeny produced per female by counting the number of 1st instar nymphs after birth. Our final sample size was 12 females for each group for the assessment of pregnancy length and the number of progeny produced.

### Milk transcription after sleep deprivation: chronic disturbance trials

To determine mechanisms for the increase in pregnancy duration, milk protein transcription levels were assessed, which have been shown in other viviparous insect systems to be directly involved in pregnancy duration (Attardo et al., 2006; Benoit et al., 2014, 2012; Denlinger and Ma, 1974; Jennings et al., 2020). As before, virgin females were mated to ensure pregnancy, and then experienced chronic sleep disturbance for 10 days, starting from 50 days into the pregnancy cycle. The disturbance consisted of removing the entire cage and vibrating for 5 minutes as previously documented (Ajayi et al. 2023). Disturbances occurred either not at all (control), twice (5 and 10 hours into the photophase period), four times (2.5, 5, 7.5 and 10 hours into the photophase), or condensed into the first 2 hours of the photophase period to allow the cockroaches a 10 hour period undisturbed in the photophase (secondary control) (Fig. 1d). Following this 10 day period of chronic disturbance, embryos were removed before the samples were frozen at −70°C and processed in biological replicates of two individuals (6-8 replicates). RNA was extracted with the use of Trizol (Invitrogen) based on the manufacturer’s protocols. Complementary DNA (cDNA) was synthesized using a cDNA Synthesis Kit (Thermo Scientific) from 1µg of RNA. KiCqStart SYBR Green qPCR ReadyMix (Sigma Aldrich) was utilized in all reactions with gene specific primer sets. Quantitative PCR was conducted in an Illumina Eco to assess milk transcript levels in biological replicates based on previously developed methods (Jennings et al., 2020). Expression levels were normalized using the ΔΔCq method as previously described (Jennings et al., 2020).

### Statistics and data analysis

All statistics were performed in R 4.3.3 (R Core Team, 2024) and most current compatible packages as of December 2024. The Shapiro test (dplyr) for normality and Breusch-Pagan test and White test (whitestrap) for heteroskedasticity were performed on each dataset, with normally distributed sets analyzed using ANOVA (dplyr) and paired T-tests (dplyr) and non-normal sets using the Kruskal Wallace (dplyr) and Dunn test (FSA). Where possible, heteroskedasticity and abnormality were corrected using a log transformation. A Bonferroni correction was used when multiple groups were compared. All statistics are reported in full in Supplemental tables 1–4. In addition to the packages associated with Rethomics (Geissmann et al., 2019): damr, behavr, ggetho, sleepr, scopr, and zeitgebr, the following R packages were utilized for data processing and visualization: ggplot2 (Wickham, 2016), viridis (Garnier et al., 2024), lmtest (Zeileis A, Hothorn T, 2002), whitestrap (Pérez JL, 2020), dplyr (Wickham et al., 2023), FSA (Ogle DH, Doll JC, Wheeler AP, Dinno A, 2023), devtools (Wickham et al., 2022), tidyverse (Wickham et al., 2019), hrbrthemes (Rudis, 2024), lattice (Sarkar, 2008), car (Fox and Weisberg, 2019), data.table (Barret, 2024), gcookbook (Chang, 2018), ggpubr (Kassambara A, 2023), and RColorBrewer (Neuwirth, 2002).

## Results

### Assessment of basic parameters of diurnal activity: Single monitor trials

Visual observation of rhythmicity and comparison of day/night averages demonstrate general nocturnal patterns for *D. punctata*. A majority of sleep occurred during photophase, and most activity occurred in the early parts of scotophase, followed by a lower activity period and declining until a substantial suppression during sunrise, when general activity was extremely low (Fig. 2a,b). The peak activity period occurred during the two hours following sunset, during which all sex/pregnancy status groups showed an increase in exploratory behavior, followed by a return to a moderate activity for the remainder of the night. No overall difference in activity levels between groups was noted (Kruskal-Wallis test: χ2=2.4867, df=2, *P* = 0.289). As expected, the population showed higher sleep during the day than at night (Kruskal–Wallis test: χ2=80.063, df=1, *P* < 0.0001), demonstrating nocturnal patterns (Fig. 2c). Sleep was significantly lower in males (Wilcox Test : W = 2502, df = 1, *P* = 0.006), with pregnant and non-pregnant females showing no significant difference (Dunn test: Z=0.983, *P* = 0.977) (Fig. 2e). This was true for both day and night periods. Inversely, beam cross data showed that males had the highest activity over all periods, and that activity trends were higher during scotophase compared to photophase (Kruskal-Wallis test: χ2=80.063, df=1, *P* << 0.0001, Wilcox test: W=14068, *P* << 0.0001). Based on these trends during this key scavenging period, we hypothesized that this may be a time when more fine-grained behavioural differences between pregnant and nonpregnant females may be observed with detailed tracking.

**Fig. 2.**
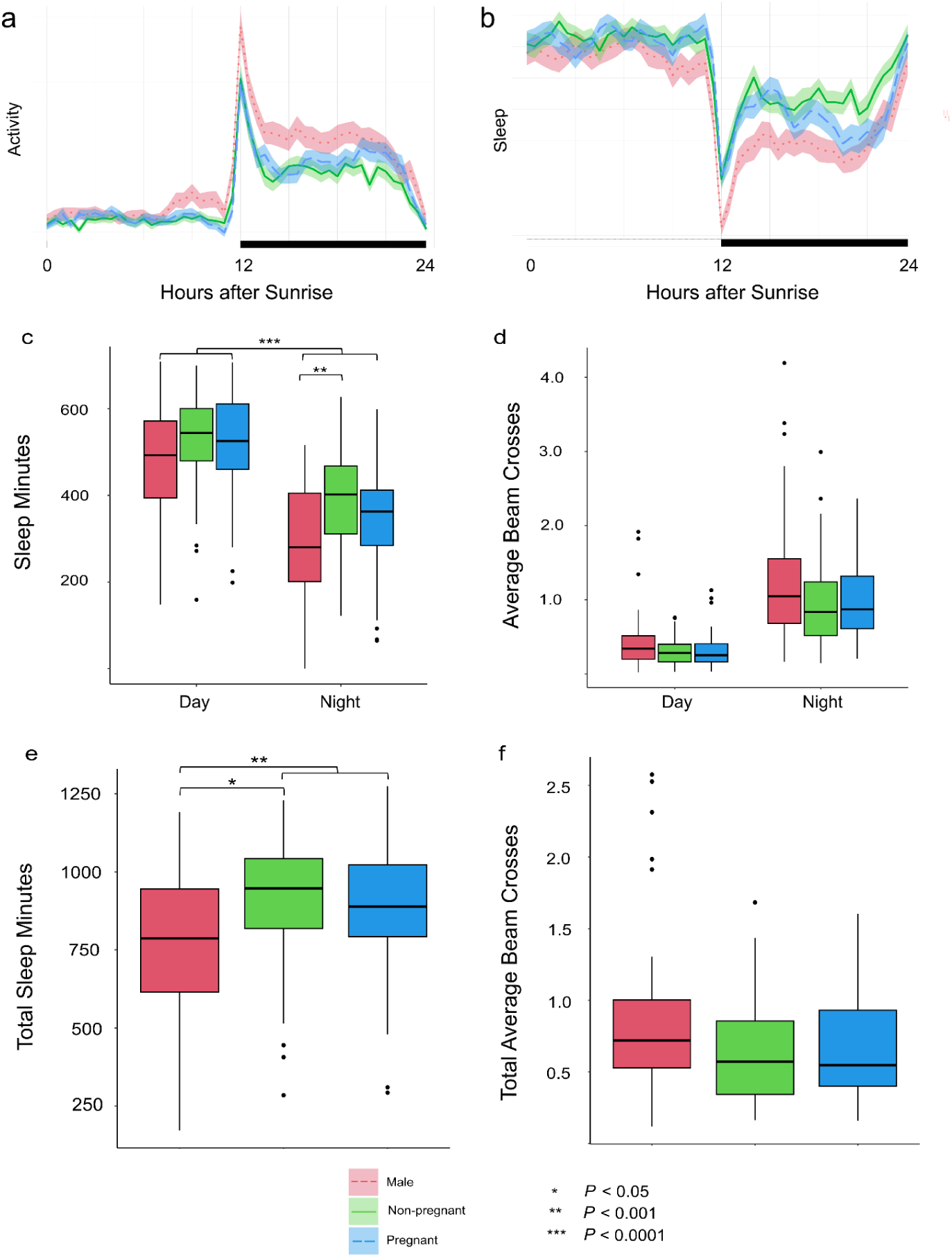
Peak in activity for *Diploptera punctata* following sunset. a. Activity levels (measured as, total beam crosses) during photophase and scotophase. We identify sunset as the key scavenging period, and activity remains moderate throughout the night, with very low activity during daylight hours. b. Sleep patterns (measured as 30 minutes without movement as the sleep threshold) show an inverse trend, high during photophase and low during scotophase, with a major dip just after the transition. C. Across all groups, individuals spend more time sleeping during the day, but d. Activity levels show no significant differences between the day and night with most activity occurring during the dusk.. Additionally, e. Total sleep minutes over the 24 hour period show significantly higher sleep in females (both pregnant and non-pregnant) compared to males, while f. Activity levels over the entire 24 hour period do not show significant differences by group. All statistics are reported in Supplemental Table 1.

### Basic assessment of cockroach emergence from harborages: dual monitor trials

Males showed higher activity patterns (measured through beam cross rate) both at short and far distances compared to females during both sunset (ANOVA: F = 6.428, df = 1, *P* = 0.0121), and night (ANOVA: F = 7.11, df = 1, *P* = 0.00836) (Fig. 3). Pregnant and non-pregnant females showed no significant difference in activity at the far sensor, indicating no difference in deep exploration related to scavenging behavior. Pregnant and non-pregnant females also showed no statistically significant difference at the close sensor; however, pregnant cockroaches did show a trend towards lower total emergence at the close sensor when compared to non-pregnant females (Fig. 3) (Dunn Test - Pregnant vs Non-pregnant: Z = 1.340, *P* = 0.541). The population showed a significantly greater emergence at the close monitor than deep exploration at the far monitor during both sunset (ANOVA: F = 33.33, df = 1, *P* = 3.34e-08) and night (ANOVA: F = 34.89, df = 1, *P* = 1.71e-08) periods. In addition, in this assay, all groups, particularly males, visually show an anticipatory rise in activity leading up to sunset before dimming begins (Fig. 3a).

**Fig 3.**
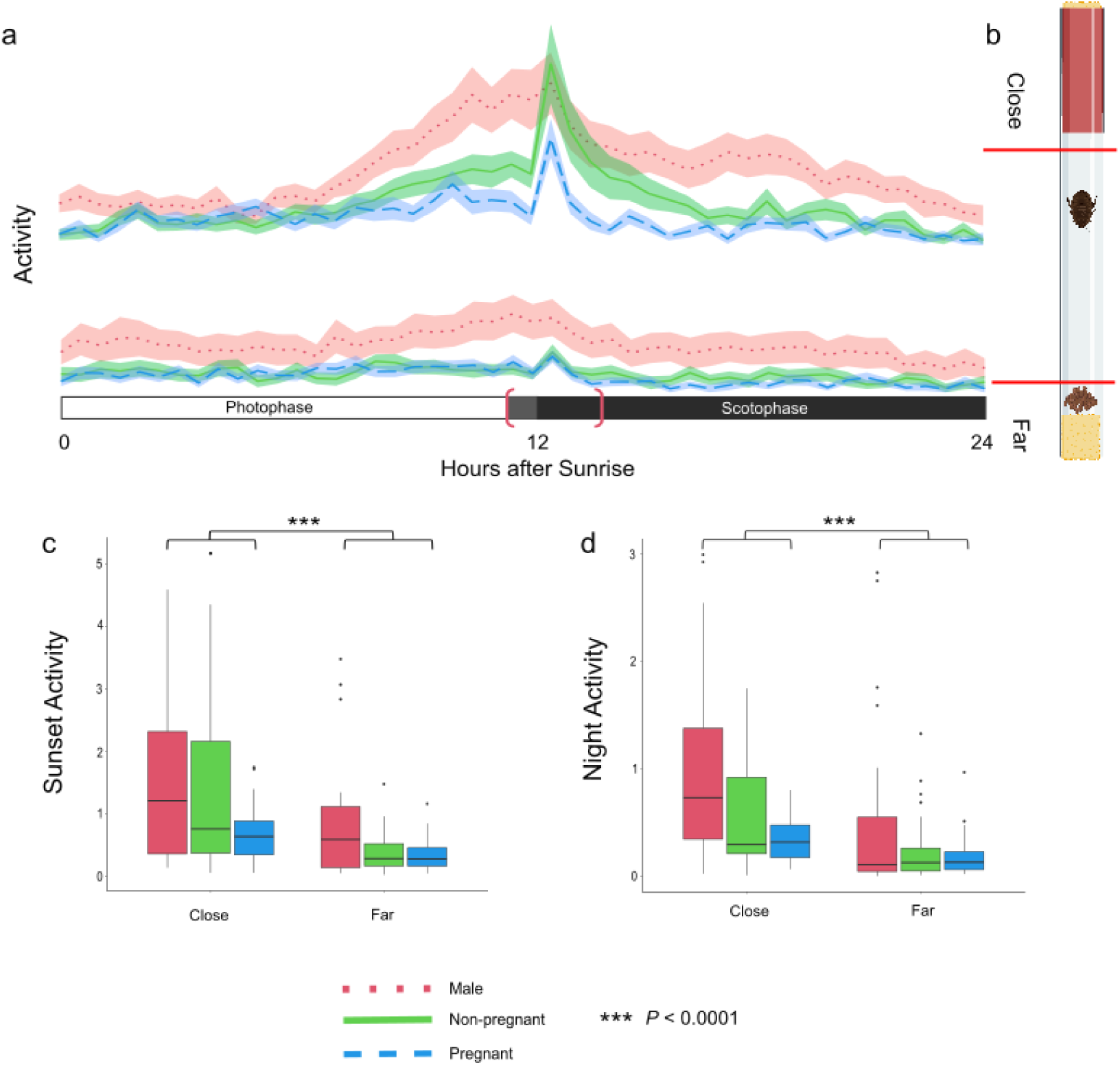
Dual monitor tracking on emergence of *Diploptera punctata* from harborages. a. Beam crosses displayed by sex/pregnancy status and position. Photophase is indicated in white and scotophase in black, with grey indicating the dimming period. For statistical purposes in this figure, ‘Sunset’ is a three hour period defined by the bracket (11:30 - 14:00). Both close and far monitor readings indicate a similar nocturnal rhythm as established in single monitor trials, including a peak in activity following sunset. At the close level, pregnant females trend towards lower emergence than males and non-pregnant females. b. Tube assay setup. Close and far infrared motion monitors are indicated by red lines. Created in BioRender. Frigard, R. (2024), BioRender.com/o34j999 c and d. Emergence trend averages are significantly higher at the ‘close’ level during both the sunset (c) and at night (d) for all groups. There is no statistically significant difference between the activity of pregnant and non-pregnant females during either sunset (c) or night (d) at either monitor, but males are significantly more active during both periods than females. All statistics are reported in Supplemental Table 2.

### Detailed analysis of harborage use and emergence at sunset: video tracking

Based on the trends from these previous trials, specific behavior patterns and differences between sex/pregnancy status groups were investigated in more detail by tracking individuals continuously over a two hour sunset period. Depth of emergence from harborage was significantly different between all three sex groups, with pregnant females showing significantly shallower emergences than other groups (Dunn Test - Male vs Pregnant: Z = - 4.014, *P* = 1.6e-2), (Dunn Test - Pregnant vs Non-pregnant: Z = 5.279, *P* = 3.9e-7). Against our expectations, non-pregnant females showed the deepest emergences, even in comparison to males (Dunn Test - Male vs Non-pregnant: Z = 2.777, *P* = 1.8e-4). The number of emergences was significantly lower for pregnant females compared to both males and non-pregnant females (Dunn Test - Male vs Pregnant: Z = 5.850, *P* = 1.5e-8), (Dunn Test - Non-pregnant vs Pregnant: Z = 3.118, *P* = 5.5e-3). There were no significant differences in the duration, but pregnant females tended to take longer trips (Fig. 4c,d).

**Fig 4.**
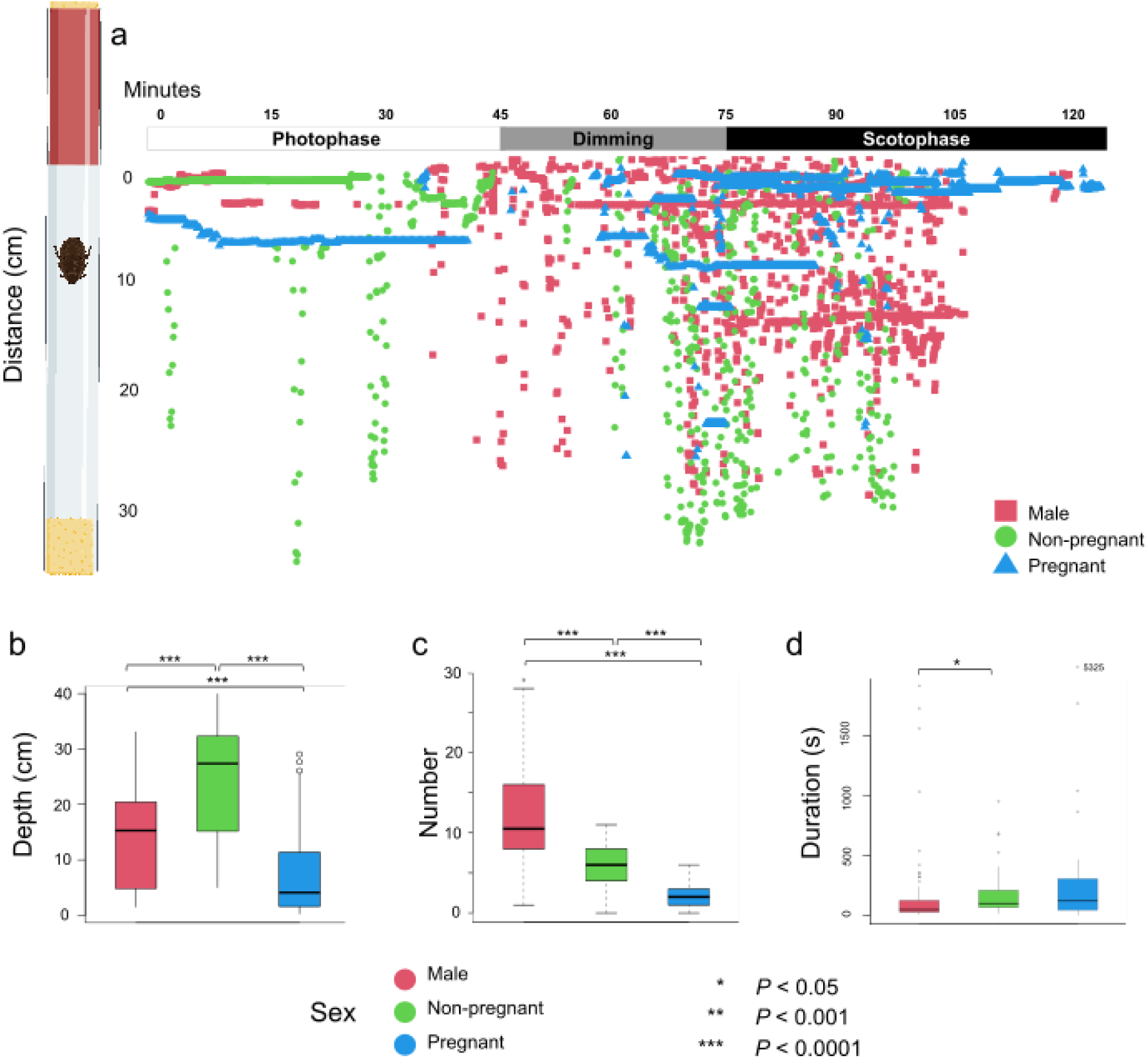
Video tracking of emergence trends of *Diploptera punctata* from harborages. **a. Emergence data from video is plotted over the two hour sunset period. Males show an immediate jump in activity at dimming onset, with females following at full darkness.** This two hour period is a portion of the 2.5 hour period described in the dual monitor trials, centered on the dimming period. Visible differences in activity patterns are present. **Tube setup diagram** included for reference. Created in BioRender. Frigard, R. (2024) BioRender.com/x73l781. **b. Depth is significantly different between all sex/pregnancy status groups**, with pregnant females showing the lowest average depth of exploration. **c. Number** of emergences is significantly different between all three groups, with males showing the most and pregnant females showing the least. **d. Durations of Emergences are not significantly different between pregnant and non-pregnant females**, but pregnant females trend towards longer emergences, and visual trends confirm this. All statistics are reported in Supplemental Table 3.

**Fig. 5.**
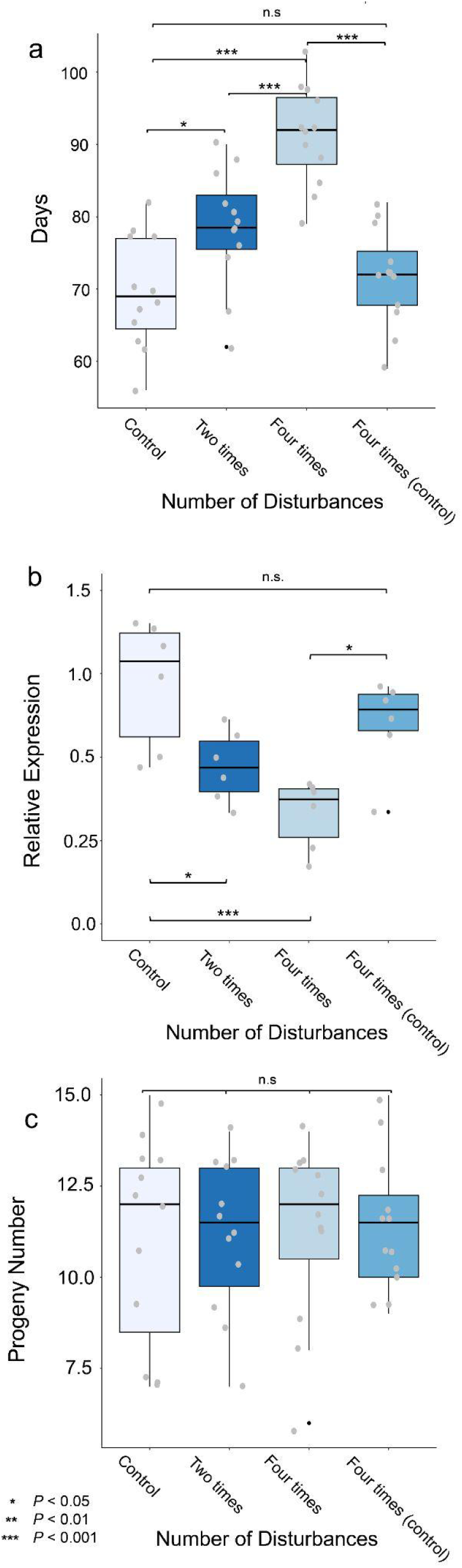
Impact of chronic sleep disturbance on pregnancy of *Diploptera punctata*. **a. Gestational period** shows a strong dose-dependent relationship between a number of disturbances during the sleeping period (photophase) and pregnancy duration. A second control group was disturbed four times during early photophase rather than throughout and showed no difference in pregnancy duration compared to the control. **b. Relative milk protein transcriptional levels** were measured in groups experiencing different chronic sleep deprivation. Again, a dose-dependent relationship was observed, with a greater number of photophase disturbances corresponding with low transcriptional levels. The second control in this trial also indicated no significant impact on milk protein transcription by chronic sleep deprivation during the nighttime active phase. **c. Progeny number** was measured and no significant difference was displayed between the groups. All statistics are reported in Supplemental Table 4.

### Pregnancy length and progeny number following sleep deprivation: chronic disturbance trials

On average, the control group of *D. punctata* (experiencing no disturbance) had a gestational length of just under 70 days. When pregnant females were chronically disturbed four times per night over a 12 hour period, pregnancy length was increased significantly by nearly 25 days (Pairwise T-test: F = 21.09, df = 2, *P* = 3.5e-08). This showed a dose-dependent relationship, with pregnant individuals only disturbed twice per cycle showing a mean increase of about 10 days (Pairwise T-test: F = 21.09, df = 2, *P* = 0.0326). A secondary control group was disturbed four times early in photophase, and no significant difference in pregnancy duration was observed.

### Milk transcription after sleep deprivation: chronic disturbance trials

The expression levels for the milk protein decreased with increasing disturbances per day. Compared to the control group (no disturbance), relative transcription levels of milk proteins decreased by nearly 50% in pregnant females disturbed four times during the day (Pairwise T-test: F = 8.647, df = 3, *P* = 0.00059), and roughly half that in those disturbed two times during the day (ANOVA: F=21.09, df=2, *P* << .0001, Pairwise T-tests: Residuals: df=44, sum sq 2409, mean sq 54.8, *P* = 0.033). A secondary control group disturbed during scotophase showed transcription levels not significantly different from the control group. No differences were noted in the number of progeny following chronic disturbance, suggesting that the decreased milk protein production is compensated by increasing the duration of the pregnancy to yield an equal number of progeny.

## Discussion

Our single monitor trials confirm *Diploptera* as a vespertine to nocturnal species (Stephenson et al., 2007), with a peak in activity in the two hours during and following sunset, and sleep occurring mainly during photophase. More detailed tracking and the inclusion of harborages in the dual monitor and video trials allowed us to consider natural hiding behaviors (Broom and Ruxton, 2005; Tobler, 1983), and develop a more robust model of *Diploptera* daily rhythms. Males are more active across the daily cycle than females regardless of pregnancy status, and pregnant females show activity patterns, such as reduced emergences, that indicate aversion to risk-taking and decreased scavenging efficiency. In addition, sleep disturbance leads to extension of gestational duration and a decrease in milk protein transcriptional levels.

The high-activity post-sunset period is when most scavenging behavior occurs and is key to nutrient consumption and hydration for *Diploptera*, as these are the main reasons for exploration in a scavenging species. For males, locating a female to mate with and engaging in territorial disputes are another major purpose of exploration, which we see reflected by the increased activity of males at all time points across experiments compared to females, as observed in other species (Carrel and Tanner, 2002). This post-sunset period is a logical time for peak activity, as many predators and parasites of cockroaches are nocturnal or matutinal (active in the early morning) and may be least active during this early evening period (Cameron, 1955).

Previous studies suggest differences between male and female patterns of activity in cockroaches; male American cockroaches show a more robust daily rhythm, while females have a more variable rhythmic activity cycle (Lipton and Sutherland, 1970). Although this previous literature shows greater variance among cockroach females, our study indicated that female *Diploptera* have a consistent light cycle dependent rhythmicity similar to other groups. This may be because oviparous species undertake a shorter and more extreme progeny development period (Roth and Willis, 1952), and that the extended pregnancy and viviparous strategy of *Diploptera* allow for more regular rhythms in scavenging and sleep.

During the dual monitor trials, we were able to investigate movement in more detail using the close and far sensors, which we continued using individual continuous video tracking. Pregnant females were less likely to leave the harborage, and when they did, spent less time scavenging on a distant food source. This may indicate increased risk aversion or energy conservation strategies leading to impaired feeding in pregnant females, which aligns with their decreased speed and mobility (Greven et al., 2014), as well as protection of the brood. While this is somewhat counter to the assumed greater nutritional requirements in pregnant females, studies in pigs indicate that pregnancy is associated with decreased activity (Marchant-Forde and Marchant, 2004), and in humans, pregnancy is associated with decreased risk-taking behavior (Chen et al., 2020). Of importance, the cockroaches used in our studies were well-fed and likely have sufficient nutrient reserves for milk production, and more scavenging may have been noted if food sources are less available or lower quality before the assays. Modeling of feeding strategies in another viviparous insect, the tsetse fly, suggests that tsetse flies are most vulnerable to predation during the early period after a bloodmeal, ergo during the early stages of their pregnancy, and that they may avoid feeding during this period (Hargrove and Williams, 1995). In addition, specific study of behavior over pregnancy and across nutritional states may indicate changing preferences as progeny develop, and elucidate more about the role of mother:offspring nutritional conflict, as well as conflict between increased risk-aversion and increased nutritional requirements. In addition, a different method may facilitate the analysis of postural data, which would allow us to see if the longer periods of stillness observed in pregnant females are true sleep-like states (Helfrich-Förster, 2018), and to investigate the possibility of a sleep rebound following deprivation as seen in mosquitos (Ajayi et al., 2022). Investigation toward the quality of sleep during pregnancy would allow us to make this distinction and explore more deeply the metabolic and hormonal changes during pregnancy which may impact circadian rhythm.

Just as daily rhythms were impacted by sex and pregnancy status, we found chronic sleep disturbance to have effects on key pregnancy factors such as duration and milk expression levels. Previous studies in mammals such as rats (Li et al., 2023) and humans (Hammer et al., 2019), demonstrate that sleep deprivation is associated with miscarriage and other complications for both mothers and young. In our model, we believe that the decrease in milk protein transcription is directly related to the elongation of the pregnancy cycle, with the increased length of time a compensatory mechanism employed by the mother to ensure progeny number. When chronic sleep disturbance occurs, milk protein levels decline, decreasing nutrients available to the embryos during development, particularly protein. *Diploptera* milk consists of 45% protein, with the remainder composed of mainly carbohydrates and lipids, and of the carbohydrates, 26% are incorporated into protein as well. In addition, the gut contents of embryos are almost entirely protein (Ingram et al., 1977). An earlier study in *Diploptera* has shown that protein deprivation led to a decrease in progeny number but little change in progeny length (Stay and Coop, 1973). This is an interesting juxtaposition to our data, which demonstrated no change in progeny number following chronic sleep deprivation. In addition, chronic sleep deprivation is known to raise the metabolic rate of male *Diploptera* (Stephenson et al., 2007). These previous studies, combined with the finding that milk protein transcription is decreased following chronic disturbance, lead us to believe that there may be a decrease in weight or other aspects of fitness in embryos produced by sleep deprived mothers. However, the extension of the gestational period could fully compensate for the decreased milk transcription, resulting in progeny of a normal weight. This pregnancy extension has been observed in viviparous insects in relation to insufficient milk production (Benoit et al., 2014; Jennings et al., 2020). Further study is required to measure these consequences of sleep deprivation for pregnancy outcomes, with particular focus on whether individual offspring are smaller as a result of the longer pregnancy duration, as well as whether there are costs to mothers of extended pregnancy duration for their own lifespan or future reproduction.

If *Diploptera* extends their gestational period, this could have implications regarding the mechanism that induces, or indeed prevents, birth in this species. The balance between extending and terminating pregnancy has been studied across many organisms. In another live-bearing species and significant disease vector, the tsetse fly, nutrient deprivation is associated with higher rates of spontaneous abortions (English et al., 2023), and in humans, lack of development and low fetal weight are associated with miscarriage (Marques et al., 2024; Poorolajal et al., 2014). In fact, postterm pregnancies are associated with lower birth weights and other delivery complications, indicating that the factors which lead to fetal malnutrition are not overcome by extended pregnancy length (Galal et al., 2012). While the exact mechanism that induces labor is unknown, in certain mammals it is thought that a mature embryo releases proteins into the maternal bloodstream (Condon et al., 2004), which, alongside other hormonal and potentially immunological changes (Liao et al., 2005), encourage labor. In banded mongooses, dominant females engage in behavior to synchronize the birth timing of all females in the group, potentially to reduce infanticide by stronger pups (Gilchrist, 2006). Uncovering whether or not a mechanism similar to mammalian birth timing strategies exists in *Diploptera*, and if the control is solely the mother or dynamics between the mother and offspring, could have implications on the mechanisms used in many other live-bearing species. The hormonal regulation of birth in *Diploptera* most likely involved a increase in JH and a decline in ecdysteroids, which yield a reduction in the synthesis of milk proteins a few days before birth and the start of the next cycle of oogenesis (Rankin and Stay, 1985; Ter Wee and Stay, 1987; Tobe et al., 1985).

In conclusion, we use detailed tracking technology to show that daily rhythms change during pregnancy in *Diploptera punctata* and chronic sleep deprivation yields negative effects on pregnancy, increasing gestational length and decreasing the transcription of milk proteins critical to embryo development. In addition, adjustments to daily rhythms and scavenging behavior are made to accommodate changes in metabolism and nutrient gathering behavior. Altogether, these results demonstrate that sleep in the Pacific beetle mimic cockroach is vital to metabolic functions necessary during pregnancy and inversely that behavior related to daily rhythms and scavenging are altered to accommodate these needs and possibly to prevent risk taking or decrease energy expenditure during pregnancy. Our work highlights the tractability of this model to investigate evolutionarily conserved associations between sleep and pregnancy, yielding insights on the mechanisms linking sleep and pregnancy outcomes across diverse taxonomic groups.

## Acknowledgments

Partial funding was provided by the National Institute of Allergy and Infectious Diseases of the National Institutes of Health under Award Number R01AI148551 and National Science Foundation DEB1654417 (to J.B.B. for shared computational and reusable equipment resources). Partial funding was provided by the UPRISE program of the University of Cincinnati (R.F.). S.E. was supported by a UKRI Future Leaders Fellowship (MR/W007711/1) and Royal Society Dorothy Hodgkin Fellowship (DH140236).

## Supplemental Material

**Supplemental Table 1.**

| **Figure 2 Statistics**<br>All groups $n = 80$ | **Variance** | **df** | **P value** |
| --- | --- | --- | --- |
| **Beam Count by Group Kruskal Wallis Test** | $\chi^2 = 2.6687$ | 2 | 0.2633 |
| **Beam Count by Phase Kruskal Wallis Test** | $\chi^2 = 2.486$ | 2 | 0.2885 |
| **Day Beam Count by Group Kruskal Wallis Test** | $\chi^2 = 2.3521$ | 2 | 0.3085 |
| **Night Beam Count by Group One-way ANOVA Test** | F = 1.035<br>Residuals: df 128, sum sq 53.93 , mean sq 0.4213 | 2 | 0.358 |
| **Total Sleep Minutes by Group**<br>Kruskal-Wallis Test<br>Dunn Test<br>Male vs Non-Pregnant<br>Male vs Pregnant<br>Non-Pregnant vs Pregnant | $\chi^2 = 8.5176$<br>Z = -2.873<br>Z = -1.851<br>Z = 0.983 | 2 | 0.01414 *<br>0.01219522 *<br>0.19246150<br>0.97740244 |
| **Sleep Minutes by Phase**<br>Kruskal-Wallis<br>Wilcox Test | $\chi^2 = 80.063$<br>W = 14068 | 1 | < 2.2e-16 ***<br>< 2.2e-16 *** |
| **Day Sleep Minutes by Group Kruskal Wallis Test** | $\chi^2 = 2.6506$ | 2 | 0.2657 |
| **Night Sleep Minutes by Group**<br>Kruskal-Wallis<br>Dunn Test<br>Male - Non-Pregnant<br>Male - Pregnant<br>Non-Pregnant - Pregnant | $\chi^2 = 10.699$<br>Z = -3.26<br>Z = -1.82<br>Z = 1.389 | 2 | 0.003342264 **<br>0.003342264 **<br>0.204006072<br>0.494620500 |
| **Sleep Minutes Male vs Female**<br>Kruskal-Wallis | $\chi^2 = 7.552$ | 1 | 0.005994 ** |
| Wilcox Test | W = 2502 |  | 0.006039 ** |
| Beam Counts Male vs Female One-way ANOVA Test | F = 2.292<br>Residuals: df 129, sum sq 49.65, mean sq 0.3849 | 1 | 0.133 |

**Supplemental Table 2.**

| **Figure 3 Statistics**<br>Male, n = 28<br>Non-pregnant, n = 33<br>Pregnant, n = 31 | **Variance** | **df** | **P value** |
| --- | --- | --- | --- |
| **Sunset, Close Monitor, by Sex group**<br>Kruskal-Wallis<br>Dunn Test<br>Male vs Non-pregnant<br>Pregnant vs Male<br>Pregnant vs Non-pregnant | $\chi^2 = 6.1981$<br>Z = 1.228<br>Z = 2.487<br>Z = 1.340 | 2 | 0.04509 *<br>0.65806883<br>0.03859808 *<br>0.54068719 |
| **ANOVA, Sunset, Far Monitor, by Sex group** | F = 1.819<br>Residuals: df 88, sum sq 97.56, mean sq 1.109 | 2 | 0.168 |
| **Sunset, Both Monitors, by Sex group**<br>ANOVA<br>Pairwise T-tests<br>Male vs Non-pregnant<br>Pregnant vs Male<br>Pregnant vs Non-pregnant | F = 3.538 | 2 | 0.0311 *<br>0.19<br>0.03 *<br>1.00 |
| **ANOVA, Sunset, by Monitor**<br>Close vs Far | F = 33.33<br>Residuals: df 180, sum sq 213.76, mean sq 1.19 | 1 | 3.34e-08 *** |
| **ANOVA, Night, by Monitor**<br>Close vs Far | F = 34.89<br>Residuals: df 180, sum sq 260.5, mean sq 1.45 | 1 | 1.71e-08 *** |
| **ANOVA, Sunset, by Sex**<br>Male vs Female (all) | F = 6.428<br>Residuals: df 180, sum sq 244.62, mean sq 1.359 | 1 | 0.0121 * |
| **ANOVA, Night, by Sex** | F = 7.11 |  |  |
| Male vs Female (all) | Residuals: df 180, sum sq 299.22, mean sq 1.662 | 1 | 0.00836 ** |

**Supplemental Table 3.**
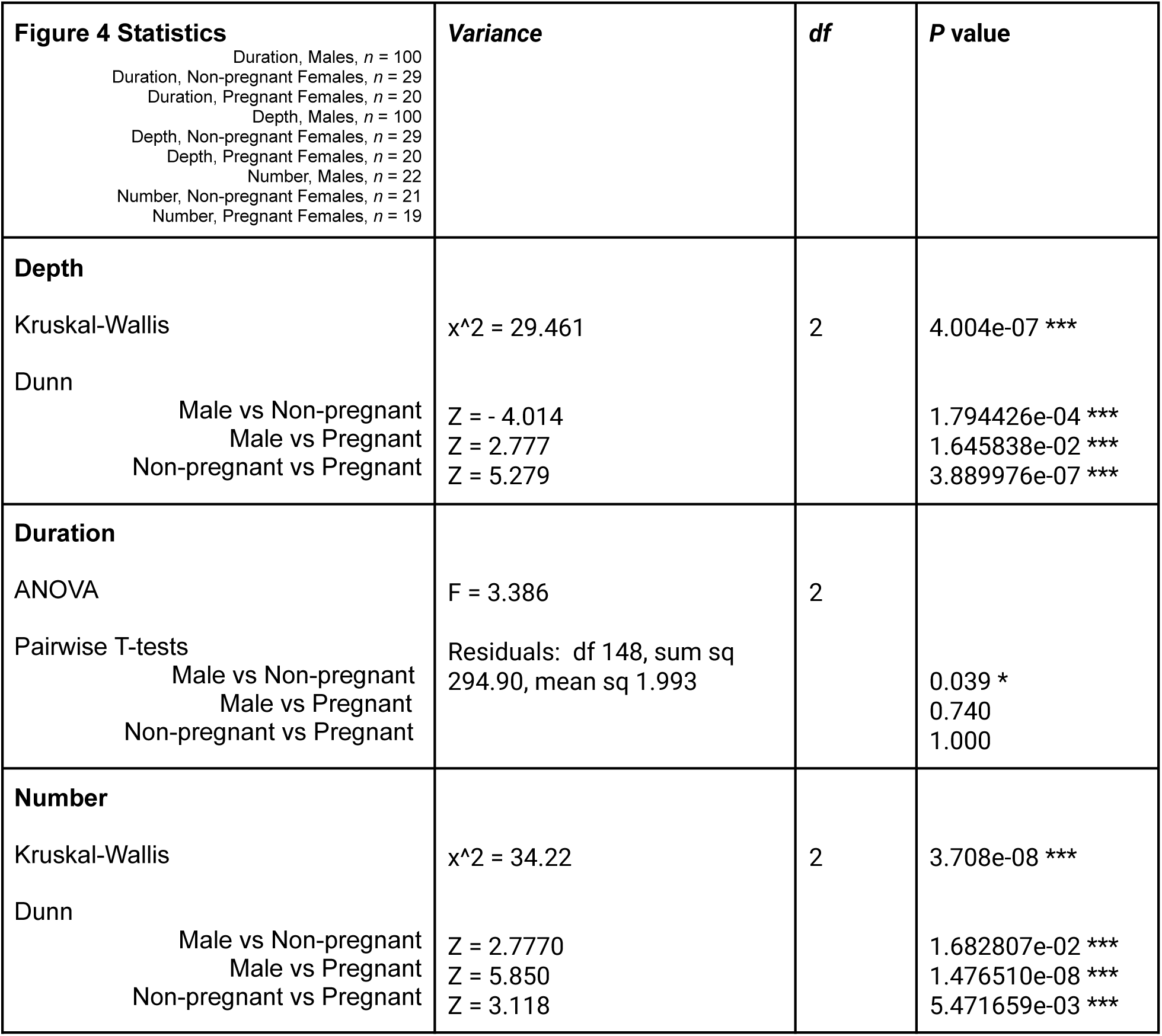

| **Figure 4 Statistics**<br>Duration, Males, $n = 100$<br>Duration, Non-pregnant Females, $n = 29$<br>Duration, Pregnant Females, $n = 20$<br>Depth, Males, $n = 100$<br>Depth, Non-pregnant Females, $n = 29$<br>Depth, Pregnant Females, $n = 20$<br>Number, Males, $n = 22$<br>Number, Non-pregnant Females, $n = 21$<br>Number, Pregnant Females, $n = 19$ | **Variance** | **df** | **P value** |
| --- | --- | --- | --- |
| **Depth**<br>Kruskal-Wallis<br>Dunn<br>Male vs Non-pregnant<br>Male vs Pregnant<br>Non-pregnant vs Pregnant | $\chi^2 = 29.461$<br>Z = -4.014<br>Z = 2.777<br>Z = 5.279 | 2 | 4.004e-07 ***<br>1.794426e-04 ***<br>1.645838e-02 ***<br>3.889976e-07 *** |
| **Duration**<br>ANOVA<br>Pairwise T-tests<br>Male vs Non-pregnant<br>Male vs Pregnant<br>Non-pregnant vs Pregnant | F = 3.386<br>Residuals: df 148, sum sq 294.90, mean sq 1.993 | 2 | 0.039 *<br>0.740<br>1.000 |
| **Number**<br>Kruskal-Wallis<br>Dunn<br>Male vs Non-pregnant<br>Male vs Pregnant<br>Non-pregnant vs Pregnant | $\chi^2 = 34.22$<br>Z = 2.7770<br>Z = 5.850<br>Z = 3.118 | 2 | 3.708e-08 ***<br>1.682807e-02 ***<br>1.476510e-08 ***<br>5.471659e-03 *** |

**Supplemental Table 4.**

| **Figure 5 Statistics**<br>All groups, Expression, $n = 6$<br>All groups, Duration, $n = 12$<br>All groups, Progeny, $n = 12$ | **Variance** | **df** | **P value** |
| --- | --- | --- | --- |
| **Duration** |  |  |  |
| ANOVA | F = 21.09 | 2 | 1.28e-08 *** |
| Pairwise T-tests | Residuals: df 44, sum sq 2409,<br>mean sq 54.8 |  |  |
| Control vs Four times |  |  | 3.5 e -8 *** |
| Control vs Four times (control) |  |  | 1.000 |
| Control vs Two times |  |  | 0.0326 * |
| Four times vs Four times (control) |  |  | 3.6 e -7 *** |
| Four times vs Two times |  |  | 0.0006 *** |
| Four times (control) vs Two times |  |  | 0.1835 |
| **Expression** |  |  |  |
| ANOVA | F = 8.647 | 3 |  |
| Pairwise T-tests | Residuals: df 20, sum sq 1.576,<br>mean sq 0.0788 |  |  |
| Control vs Four times |  |  | 0.00059 *** |
| Control vs Four times (control) |  |  | 0.87749 |
| Control vs Two times |  |  | 0.03514 * |
| Four times vs Four times (control) |  |  | 0.02005 * |
| Four times vs Two times |  |  | 0.56465 |
| Four times (control) vs Two times |  |  | 0.78978 |
| **Progeny** |  |  |  |
| ANOVA | F = 0.071 | 3 |  |
| Pairwise T-tests | Residuals: df 44, sum sq<br>241.83, mean sq 5.496 |  |  |
| Control vs Four times |  |  | 1.000 |
| Control vs Four times (control) |  |  | 1.000 |
| Control vs Two times |  |  | 1.000 |
| Four times vs Four times (control) |  |  | 1.000 |
| Four times vs Two times |  |  | 1.000 |
| Four times (control) vs Two times |  |  | 1.000 |

